# Quark enables semi-reference-based compression of RNA-seq data

**DOI:** 10.1101/085878

**Authors:** Hirak Sarkar, Rob Patro

## Abstract

**Motivation:** The past decade has seen an exponential increase in biological sequencing capacity, and there has been a simultaneous effort to help organize and archive some of the vast quantities of sequencing data that are being generated. While these developments are tremendous from the perspective of maximizing the scientific utility of available data, they come with heavy costs. The storage and transmission of such vast amounts of sequencing data is expensive.

**Results:** We present `Quark`, a semi-reference-based compression tool designed for RNA-seq data. `Quark` makes use of a reference sequence when encoding reads, but produces a representation that can be decoded independently, without the need for a reference. This allows `Quark` to achieve markedly better compression rates than existing reference-free schemes, while still relieving the burden of assuming a specific, shared reference sequence between the encoder and decoder. We demonstrate that `Quark` achieves state-of-the-art compression rates, and that, typically, only a small fraction of the reference sequence must be encoded along with the reads to allow reference-free decompression.

**Availability:** `Quark` is implemented in `C++11`, and is available under a GPLv3 license at www.github.com/COMBINE-lab/quark.

## 1 Introduction

Compression of high-throughput sequencing reads becomes crucial with the lowering cost of sequencing technology. The rapid technological development enables the generation of petabytes of data on servers worldwide. Apart from size, often succinct representation (Pritt and Langmead, 2016) can yield very similar results with a much smaller memory footprint. Before using state-of-the-art, off-the-shelf compression tools, there is always a scope of substantial improvement as we have shown in this paper regarding rearrangement and encoding of raw read sequences. In one hand, such encoding can exploit the redundancy of highly repeated sequences from the read. On the other hand, we demonstrate that quasi-mapping, a recently-introduced proxy for traditional alignment (Srivastava *et al.*, 2016), enables selective compression of read sequences with respect to the reference sequence.

We present `Quark`, a compression method specifically designed for high throughput RNA-seq reads. On a conceptual level it introduces the idea of semi-reference-based compression, where reference sequence is used at the encoding end, but is not required for decompression. This allows `Quark` to obtain markedly better compression rates than completely reference-free tools while also eliminating the need for the encoder and decoder to share the same exact reference sequence, which also mitigates the potentially brittle dependence of a reference-based encoder on a specific reference sequence. Specifically, using quasi-mapping (Srivastava *et al.*, 2016), `Quark` locates regions of interest in the reference that are specific to the particular RNA-seq experiment being compressed, and stores only these regions for use during decoding. `Quark` is focused on sequence compression, and hence, does not currently provide a mechanism for storing the header and quality information associated with each read. Although there are very efficient approaches to these problems (Zhou *et al.*, 2014), (Malysa *et al.*, 2015), (Janin *et al.*, 2013) that could easily be coupled with `Quark`. Apart from reducing the total size of `fastq` files, the other motivation is that many tools (e.g., state-of-the-art quantification tools such as `Sailfish` (Patro *et al.*, 2014), `Salmon` (Patro *et al.*, 2016) and `kallisto` (Bray *et al.*, 2016)) do not make use of this information from the `fastq` files. In fact, the link between `Quark` and transcript quantification methods goes even deeper, as `Quark`’s notion of islands Section 3.1 naturally extends and refines the notion of fragment equivalence classes first introduced in `mmseq` (Turro *et al.*, 2011), and subsequently adopted by recent lightweight quantification approaches such as `Sailfish`, Salmon, and `kallisto`.

`Quark` is the first reference-asymmetric compression methodology of which we are aware (i.e., in terms of only requiring the reference for encoding). Further, it develops certain key connections between the redundant representation of sequence information, in terms of read compression, and the use of related ideas in the efficient likelihood factorization that has been integral to the development of fast quantification methodologies. Our analysis also provides some insights into the typical coverage / usage of unique sequence in specific RNA-seq experiments (Section 3.1). The idea of semi-reference-based compression appears very effective at improving sequence compression rates, but not limited only to RNA-seq data. Rather, we believe that the ideas we present here can be extended to genomic data and, possibly, even to long read compression.

## 2 Related Work

Given the volume of high-throughput sequencing data, there has been a substantial research focus on methods for its compression. Nucleotide sequence compression can largely be divided into two paradigms — reference-based and reference-free. In reference-based compression, a reference sequence (genome, transcriptome, etc.) must be shared by both the encoder and the decoder (Cánovas *et al.*, 2014), (Fritz *et al.*, 2011), (Li *et al.*, 2014). Alternatively, in reference-free compression (e.g. (Adjeroh *et al.*, 2002), (Bonfield, 2014), (Hach *et al.*, 2012), (Patro and Kingsford, 2015)), read sequences are compressed independent of reference sequence. This eliminates the burden of requiring the encoder and decoder to share a reference, but typically results in lower compression rates than reference-based compression.

Reference-based compressors typically start with BAM files produced by aligners such as `Bowtie 2` (Langmead and Salzberg, 2012), `bwa` (Li, 2013), and `STAR` (Dobin *et al.*, 2013). One potential bottleneck of reference-based compression is to that many approaches store a substantial amount of meta-information from the BAM file which can be regenerated by re-aligning the reads to the provided reference sequence. This problem can be easily solved storing the edits after aligning the sequences to reference. Fritz *et al.* (2011) introduced one such widely used tool `mzip` which can store alignments permitting users to avoid compressing quality score and unaligned sequences. `fastqz` (Bonfield and Mahoney, 2013) bypassed the problem of storing meta data by implementing a new alignment technique.

The recently-published CORA (Yorukoglu *et al.*, 2016) is a compressive read mapper. The working principle of CORA is interesting and worth mentioning as the concept of equivalence classes (discussed in Section 3) is also used there, but carries a different meaning. CORA converts the fastq reads to non-redundant k-mer sets. Furthermore, CORA attempts to capture redundancy within the reference genome by performing self-mapping. The positions where a k-mer maps form an equivalence class. Two equivalence classes are regarded as concordant if all positions of one equivalence class are in one nucleotide shift distance from another equivalence class.

While most existing tools fit the categorizations of reference-based or reference-free well, some methods, conceptually, lie in the middle. Quip (Jones *et al.*, 2012) performs a lightweight assembly from a small subset of the input data, and then encodes the reads with respect to this assembly, capitalizing on ideas that have been used in both reference-free and reference-based compression. `kpath` (Kingsford and Patro, 2015) is a tool that assumes a k-mer distribution given by a reference transcriptome, and compresses the reads by grouping them together by their starting k-mers and encoding them as arithmetically coded paths in a de Bruijn graph. Though this is technically a reference-based method, it is designed to encode raw sequencing reads rather than alignments, and the authors demonstrate that the performance degrades gracefully even when a substantially different reference transcriptome is used to encode a sample. The reference-free compression tool, LEON (Benoit *et al.*, 2015), constructs a probabilistic de Bruijn graph from the k-mer count table built on the reads. Reads are then mapped to the newly constructed de Bruijn graph, and are stored in the form of an anchor address, the read size, and bifurcation list. This method shares some conceptual similarity with Quip, but the fact that it builds a (probabilistic) de Bruijn graph from all of the input data allows it to typically achieve much greater compression ratios.

In this paper we present a compression tool, `Quark`, which we categorize as a semi-reference-based compressor. `Quark` takes as input raw reads and a reference transcriptome, and produces an encoding that can be decompressed without knowledge of the original reference sequence. Thus, the reference sequence is required at the encoder, but not at the decoder. The asymmetry of `Quark` is designed to match the typical asymmetry in terms of data availability and processing power between the encoder and decoder, with encoding requiring more data and computation than decoding. This semi-reference-based scheme has a number of benefits. First, making use of the reference at the encoding end allows `Quark` to obtain substantially better compression ratios than existing reference-free tools. This results in smaller files that consume less space on disk, and which are faster to transfer from e.g., a large centralized repository to a remote machine for analysis. Second, the reference-free nature of `Quark`’s decompression relieves the assumption of shared knowledge between the encoder and decoder. In data that is compressed for potentially long-term storage, it is likely that the reference used for encoding will no longer be current or in wide circulation and that the decoder will have to *re-fetch* this reference sequence. However, since `Quark` allows the decoder to recover the original data without sharing the specific reference used for encoding, it is immune to the fact that reference sequences are commonly augmented or updated.

`Quark` uses the recently-introduced concept of quasi-mapping (Srivastava *et al.*, 2016) to quickly *map* sequencing reads to a target transcriptome. The mapping information is then utilized to represent the reads. The motivation for such a scheme comes from the observation of the fact that a small fraction of the transcriptome is often sufficient to represent the vast majority of mapping reads in an experiment, and therefore, storing the entire transcriptome is often unnecessary. A decoder on the other end takes the small subset of the transcriptome sequence, that we refer to as islands, and the compressed reads to produce the uncompressed data. An overview of `Quark` is given in Figure 1. The method section is divided into three parts, in the heart of `Quark` quasi-mapping maps the raw reads to the indexed transcriptome and reports the position of the read (anchored by a series of right-maximal exact matches). In the next step, we collect the position and the target transcripts, and subsequently merge the reference sequences overlapping the mappings to yield a set of islands. Finally, the encoded read and reference sequence islands are post-processed with an off-the-shelf compression tool (here, we use `plzip`).

**Fig. 1:**
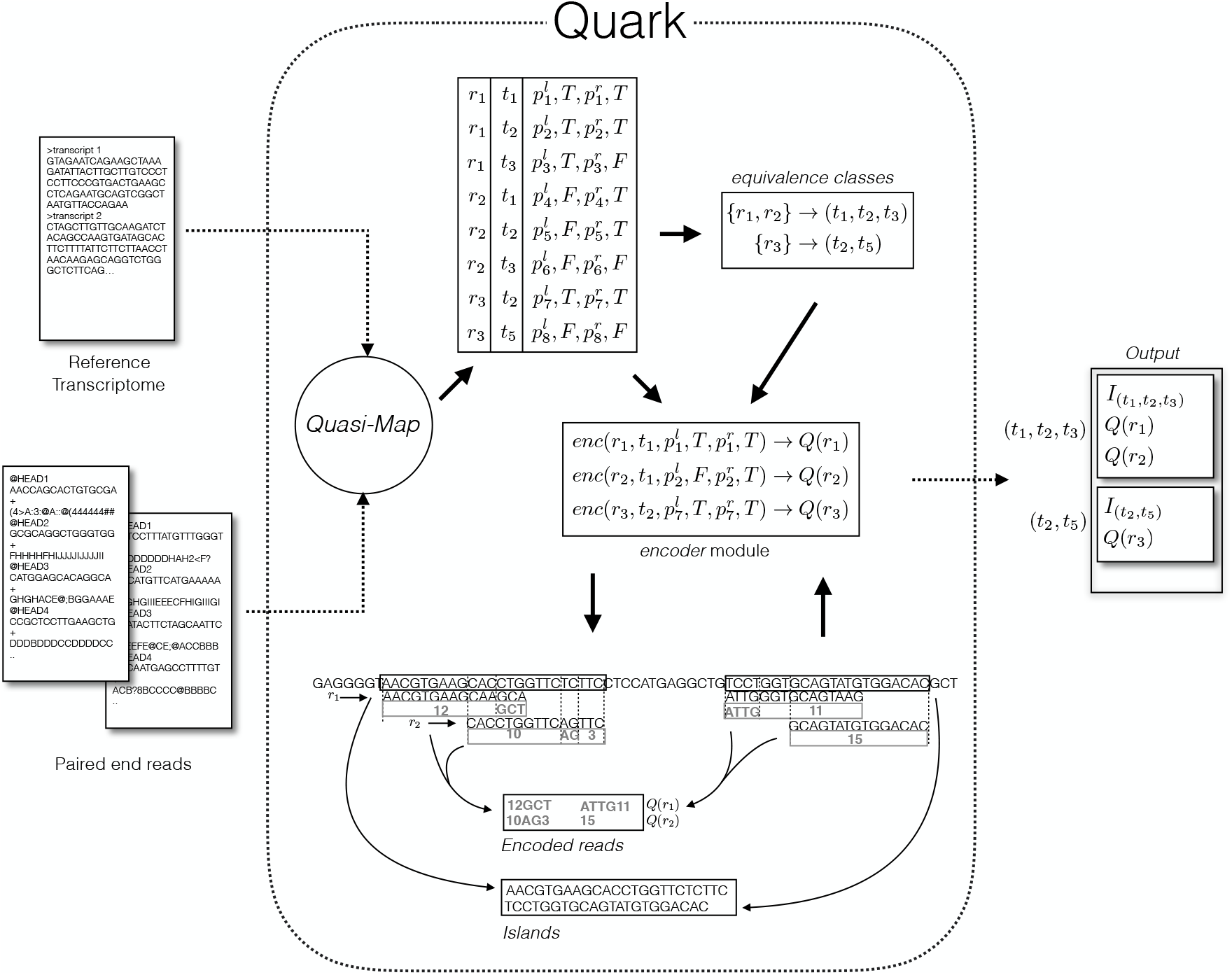
Quark uses the core component of `RapMap` which is quasi-mapping. It is used to produce a set of tuples for a paired-end read *r*_*i*_, where each tuple can be represented as (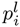, 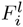, 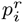, 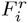). The table contaning the tuples for each read can be summarized to a set of equivalence classes as discussed in Section 3. The encoding function *Q* is explained above with two paired end reads *r*_1_ and *r*_2_. For left end of *r*_1_, there are 12 matches followed by unmatched characters. For the right end of *r*_1_, first 4 characters differ from the reference, followed by 11 exact matches, the left and right end together can be encoded as *Q*(*r*_1_) = {12GC,ATTG11}. The relevant intervals are subsequently stored as islands.

## 3 Method

`Quark` implements a semi-reference-based compression algorithm, where the reference is required for compression but not for decompression. To remove this dependency at the decoder, `Quark` encodes and stores only parts of underlying reference which are required for decompression. Thus, the output of `Quark` is self-contained in the sense that the raw reads can be recovered from the `Quark` output without the aid of any additional file. To be precise, given the reference sequence and the reads, `Quark` generates three files, *read.quark, offsets.quark* and *islands.quark.* Before describing the core algorithm of `Quark` in detail, we briefly describe the quasi-mapping concept, and the algorithm to efficiently compute quasi-mappings introduced in Srivastava *et al.* (2016), which is an integral part of the `Quark` algorithm.

Given some reference (e.g., a transcriptome), quasi-mappings identify each read with some set (possibly empty) of target sequences (e.g., transcripts), positions and orientations with which the read shares a consistent collection of right-maximal exact matches. The quasi-mapping algorithm described in Srivastava *et al.* (2016) constructs a suffix array-based index over the reference sequence. The mapping process starts with a matched k-mer that is shared between a set of transcripts and the read. If such a match exists, the algorithm tries to extend the match further by searching the interval of the suffix array for a maximum mappable prefix. The match is used to determine the *next informative position* in the reference sequences, and the same mapping process continues from that point within the query. These exact matched sequences play an important role in achieving superior compression rate of `Quark`.

On the basis of the result provided by the quasi-mapping algorithm described above, we can divide the input reads (assumed, for simplicity, to be paired-end reads) into three categories,

- *Mapped reads:* If both reads of the pair are mapped, this is an ideal situation where we can encode both ends of the read efficiently since each of the reads shares some sequence with the reference.
- *Orphan reads:* For reads in this category, we can not map both ends of the pair to the same target. In `Quark`, we encode the unmapped end of the read by writing its (encoded) sequence directly.
- *Unmapped reads:* There is no mapping at all for the read, as determined by the algorithm described above, and so the read is instead encoded using a reference-free approach. In `Quark`, un-mappable reads are encoded using the reference-free compression tool. `Mince` (Patro and Kingsford, 2015).

`Quark` follows a hierarchical approach for compression, where the mapped reads are distributed into equivalence classes according to the transcripts to which they map, and then sorted, within each class, by their starting position. Reads within the same equivalence class are very likely to share overlapping reference sequence, and hence, to be similar to each other. The encoding scheme itself is straight forward. Given the position and reference sequence, `Quark` does a linear search on the reference sequence to find the matching sequence between the read and the reference at the specified position. Though it is guaranteed to yield a match of at least *k* nucleotides if given the k-mer criterion of the mapping algorithm, typically the collection of matches covers a large fraction of the read.

`Quark`’*s read encoding*. The encoding phase of `Quark` starts with the output produced by quasi-mapping. Given a read *r*_*i*_ mapped to the reference transcriptome, `Quark` produces a tuple 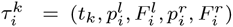 (See Figure 1), where *t*_*k*_ is the transcript sequence where the read maps, 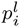 is the position where the left end maps and 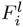 is a flag which is false if the the read has to be reverse complemented to map and true otherwise. Likewise, 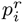 and 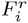 represents the corresponding position and flag for the right end of the read. When a read maps to multiple transcripts, a tuple is returned for each transcript to which the read maps. It should be noted that there are other flags that are maintained internally by quasi-mapping to keep track of orphan reads and other mapping information not currently used by `Quark`. Once the tuples τ_1_, τ_2_, for a read are obtained, they are used to place each read into an equivalence class based on the transcripts to which they map, and all reads mapping to precisely the same set of transcripts will be placed into the same equivalence class. This notion of equivalence classes has been used in the transcript quantification literature for some time (Turro *et al.*, 2011), and is described in more detail in Srivastava *et al.* (2016). As shown in Figure 1, given the mapping information for reads *r*_1_, *r*_2_ and *r*_3_, we can extract the corresponding transcripts as follows, *r*_1_ → (*t*_1_,*t*_2_,*t*_3_)*, r*_2_ → (*t*_1_, *t*_2_, *t*_3_) and *r*_3_ → (*t*_2_, *t*_5_). Intuitively, we expect that *r*_1_ and *r*_2_ are more likely to share overlapping reference sequence with each other than with *r*_3_. In addition to the transcript labels, `Quark` also associates a collection of nucleotide sequences (i.e. reference sequence to which the read maps) with each equivalence class.

The core encoding process operates on each equivalence classes individually. Given *t*_j_, a reference sequence for transcript *j*, the quasi-mapping information τ_*i*_ = (*t*_*j*_,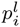,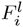,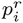, 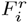) for a read, and an encoding function *Q*, `Quark` proceeds as follows. For the left end `Quark` starts a simultaneous linear scan for matches from *t*_*j*_ [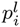] and *r*_*i*_[0], i.e., the start of the read sequence. As discussed previously, it is guaranteed that if quasi-mapping yields a mapping for this read, then the search will also find a at least a match of length k. In `Quark`, both ends of a paired end read are compressed simultaneously, and an analogous encoding procedure is used for the right end of the read. We can formalize *Q* as *Q*: Σ → S ∪ {0 — 9}, where Σ = {*A, T, G, C, N*}. Integers values are required to represent the number of matched characters. To make the `Quark` encoding more efficient, we use a four bit encoding scheme to represent each encoded read. Four bits are the minimal number of bits sufficient to represent the symbol alphabet we use for encoding because our alphabet set on the range of function *Q* is 15. The encoding scheme itself is very direct, if we have a mismatch at the start, then we put a 0 in front of the number to signify what follows is not the number of matched characters, rather it is the position where the next match characters begin.

There are two reasons why this fast and simple encoding scheme tends to yield excellent compression results. First, in NGS sequencing the raw sequences tend to contain substitution rather than indel errors. Therefore, a simple linear matching might reveal as much shared sequence as an optimal alignment. Second, even if indels or larger-scale variation are present (e.g. due to genomic divergence between the reference and sample being mapped), the repetitive reads share a common subsequence with reference, and share common edits with respect to each other, which the downstream compression algorithm can exploit.

After generating the encoded reads, `Quark` sorts each read pair by the left read’s starting position on the reference. In this way, reads that are sampled from nearby loci in the reference are placed into close proximity in the file. This step helps the downstream, off-the-shelf compressor to further efficiently compress the encoded strings. To retain the paired end read order, the right read follows the same ordering as its left mate. The read encoding process is accompanied by an island generation process, which stores the part of reference relevant for the decoding process. `Quark` also generates a separate `offset` file that contains the position of the reads.

### 3.1 Island Construction

For the purposes of compression, `Quark` makes use of islands of reference sequence that overlap the mapped reads. We define an island as a contiguous substring of some reference sequence that is completely covered by at least one read (i.e., some read overlaps each nucleotide in this substring). For each read *r*_*i*_ and its corresponding tuple τ_*i*_ = (t_k_, 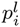, 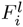, 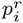, 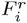), we construct an additional list *I*_i_ of intervals containing {(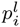, 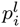 + *len*(*r*_*i*_)), (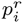, 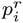 − len(*r*_*i*_))}, where *len* (*r*_*i*_) represents the length of the read. Some care must be taken to properly handle boundary conditions, which can result in situations where a read *overhangs* the beginning or end of a reference sequence. Repeating this step for each read within an equivalence class `Quark` constructs a set

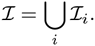

Given that intervals on the reference sequence might share some nucleotides (i.e., overlap), `Quark` merges the intervals by taking the union of the nucleotides they contain, to form maximal disjoint islands. Construction of islands from intervals is straightforward. `Quark` sorts the intervals with respect to their start positions, and a linear scan through intervals suffices to find the overlaps and merge the islands into disjoint subsets.

The use of islands aids the compression abilities of `Quark`, and additionally makes the resulting compression file self-contained, eliminating the need to assume the decoder has access to the same reference. `Quark` can identify shared regions between transcripts by the use of quasi-mapping. This further enables it to exploit redundancy and store only one island for each cluster of reads that share some nucleotide (see Figure 2).

**Fig. 2:**
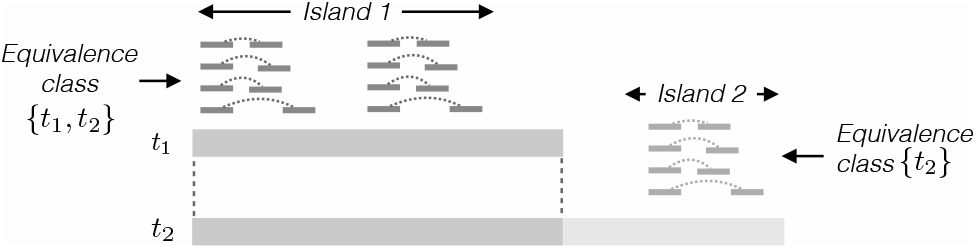
*t*_1_ and *t*_2_ are two transcripts that share a sequence. Reads labeled in dark grey are mapped into the shared region where as the reads labelled with light grey are only mapped exclusively to transcript *t*_2_, leading to formation of two equivalence classes.

The island generation process removes the redundancy of nucleotides from transcripts that share some region. Figure 2 illustrates such a situation, where a large portion of nucleotides from transcript *t*_2_ will be omitted (i.e. not included in any island). Here the reads in equivalence class {*t*_1_, *t*_2_} are completely accounted for by the island formed by the sequence from *t*_1_, so that an island corresponding to the prefix of *t*_2_ is redundant and need not be generated. However the reads in equivalence class {*t*_2_} mapped to a disjoint transcriptomic region that is not present in *t*_1_. The final set of islands (island 1 and island 2 in Figure 2) will thus contain only one representative for the redundant sequence shared by *t*_1_ and *t*_2_, so that the majority of *t*_2_ won’t be used in island creation. We further note here that this process of discovering and removing redundancy in the stored sequence from the underlying transcriptome is completely free of reference annotations, so that `Quark` works equally well when compressing the reads with respect to a *de novo* transcriptome assembly.

To study the effectiveness of constructing islands, we considred dataset `SRR635193`. After mapping to the Gencode reference transcriptome for human (version 19), we observe that out of 95, 309 transcripts, only 49, 589 transcripts are used by `Quark`. Moreover, as shown in Figure 3, there are many transcripts in which only a small fraction of nucleotides participate in islands. It is to be noted that the absence of a transcript’s sequence in islands does not necessarily imply low abundance of that transcript.

**Fig. 3:**
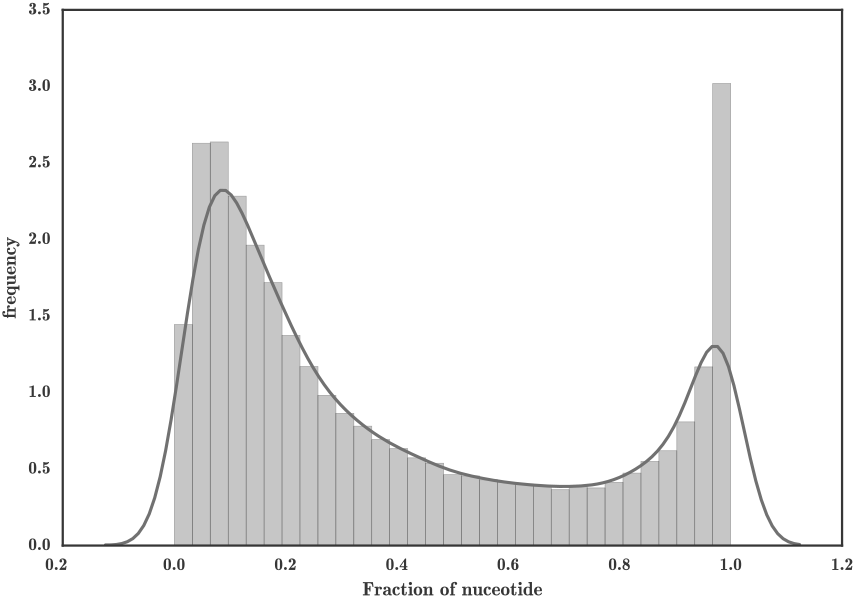
Islands consist of only a fraction of nucleotides of expressed transcripts

### 3.2 Post-processing

In addition to yielding a reduced representation of the reads, the work done by `Quark` organizes the encoded reads in a format and order that is amenable to further compression by traditional mechanisms (e.g. using programs such as `gzip, bzip2, lzip`). Given the encoding size benefits of `lzip` described by Patro and Kingsford (2015), we further process the `Quark` encodings, using `lzip` to compress the *read.quark, offsets.quark* and *islands.quark.* files, which contain the encoded reads, their offsets and the sequences of the islands, respectively. As the size of the unmapped reads can not be improved by taking advantage of quasi-mapping, we use the pre-existing *de novo* compression tool `Mince` (Patro and Kingsford, 2015) to compress these reads.

## 4 Results

Since `Quark` encodes raw sequencing reads rather than alignments, and since, from the perspective of decoding, `Quark` is reference-free, we have compared it against other *de novo* or reference-free compression tools. `Quark` does not preserve the header or quality scores from the `FASTQ` files. Therefore, we construct our baseline by extracting the read nucleotide sequences from the `FASTQ` files. As discussed in 2 we compared `Quark` with three other state-of-the-art compression tools; `LEON, SCALCE` and `Mince`. To achieve a fair comparison, we have only used the size of compressed sequences ignoring the meta information.

We used `LEON` version 1.0. `LEON` is invoked with the `- seq-only` flag to ignore the header and quality information. For `SCALCE`, version 2.8 was used. Paired-end reads are encoded with the `-r` option in `SCALCE`. Additionally we have used `-p 100` option to allow *maximally-lossy* quality compression. Unfortunately, in `SCALCE` it is not possible to completely discard the quality scores (i.e. the `. scalceq` files), since special quality values are used to encode the locations of `N` nucleotides within the reads. For the sake of completeness, column `SCALCE*`, in Table 1, records the `SCALCE` compressed file sizes, discarding the quality file altogether. This makes the file no longer decompressible, but provides a generous lower bound on how small the sequence file could get (e.g., if a different encoding scheme were used to encode the positions of “N” nucleotides in the reads). Finally we used `Mince` version 0.6. `LEON` does not provide any special way to handle paired-end reads. To allow `LEON` to obtain a maximal compression rate on paired-end data (i.e., by exploiting redundancy between both mate files), it was run on the concatenation of the left and right read pairs. As it does not alter the internal ordering of contents in the file, it is still possible to recover the original sequences, and restore the pairing, from the decoded file. We observe that this approach of running `LEON` led to slightly improved compression rates over running the tool on both read files independently (data not shown).

**Table 1.** Size of the read files after compression along with raw sequence size (bytes) PE: Paired End, SE: Single End

|  | Organism | Library Type | Raw-Sequence | Quark | LEON | SCALCE | SCALCE* | Mince |
| --- | --- | --- | --- | --- | --- | --- | --- | --- |
| SRR689233 | Mouse | PE | 2,986,245,990 | **99,028,458** | 247,687,471 | 561,022,336 | 233,134,542 | 176,947,631 |
| SRR445718 | Human | SE | 3,327,310,165 | **146,495,063** | 328,857,885 | 581,694,785 | 252,450,288 | 162,969,503 |
| SRR490961 | Human | SE | 4,961,894,468 | **159,074,135** | 466,517,518 | 745,095,275 | 299,983,580 | 174,583,153 |
| SRR635193 | Human | PE | 2,999,246,910 | **149,197,655** | 361,795,379 | 446,853,627 | 293,842,354 | 240,274,889 |
| SRR037452 | Human | SE | 421,663,860 | 54,691,651 | 90,774,360 | 111,469,817 | 66,630,751 | **54,302,557** |
| SRR1294122 | Human | SE | 3,384,221,798 | **180,413,521** | 437,077,070 | 662,252,794 | 298,757,892 | 204,850,132 |
| SRR1265495 | Human | PE | 3,432,947,700 | **158,635,313** | 305,101,315 | 709,315,147 | 320,744,415 | 242,176,319 |
| SRR1265496 | Human | PE | 2,985,375,780 | **152,612,910** | 280,749,535 | 636,872,285 | 299,286,153 | 224,755,144 |

The comparison has been carried out on eight different real datasets, as described in Table 1. There are seven fastq files from human and one from mouse (*Mus musculus*). To have a broad comparison, we have used a mixed collection of four paired end and four single end data sets.

Before discussing the enhancement in compression ratio achieved by `Quark`, we note that the compression ratios achieved by `Quark` depend on the mappability of the reads, since `Quark` uses the reference (on the encoding end) to organize the reads and place similar reads together. As shown in Figure 4, the fraction of the compressed files required to store the mappable reads is always less (usually substantially less) than the fraction of the input files consisting of mappable reads. Put differently, this demonstrates that the semi-reference-based mapping, and the encoding scheme adopted by `Quark`, is generally much more efficient than even the best of the reference-free tools. Further, it suggests that the compression achieved by `Quark` could be even further improved by assembling and mapping reads to transcripts that might not appear in the initial annotation (e.g., novel spliceforms, or tissue-specific transcripts that may not be part of the originally considered annotation). Currently, we are using the existing tool, `Mince`, to compress the unmapped reads. Although the idea of islands can be extended to the assembly of novel transcripts.

**Fig. 4:**
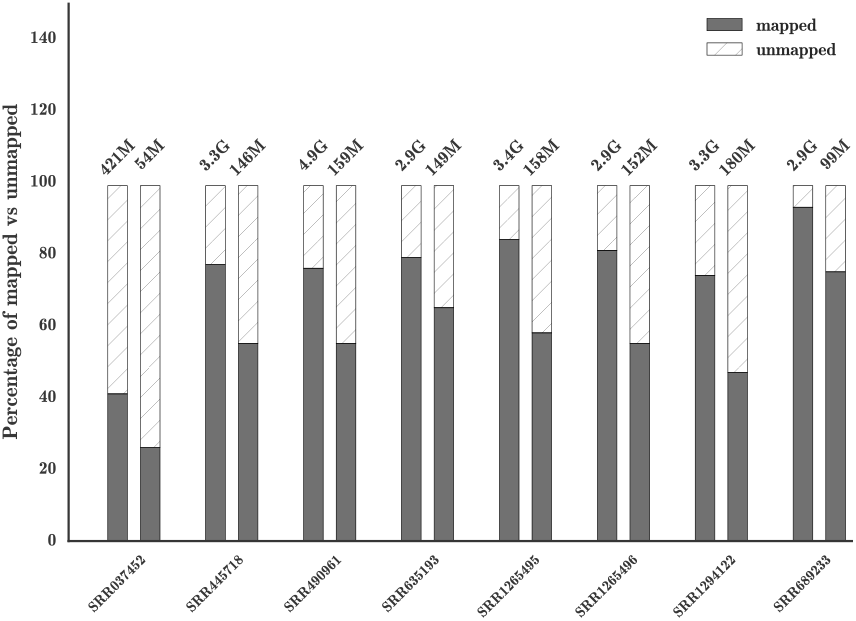
There are two bar plots corresponding to each experiment. The bar represents a normalized view of the sizes, here the grey region represents the mapped reads, whereas the striped region represents the unmapped reads. The original size of the uncompressed and compressed files are written at the top of the bar. In all cases the rate of compression for mapped reads are better than that of unmapped reads.

From Table 1, we observe that `Quark` achieves superior compression in almost all of the datasets. Additionally, for paired-end reads, the compression rate of `Quark` with comparison to the other re-ordering based methods (`SCALCE` and `Mince`) is much better. One reason for this is that, intuitively, the ordering induced by `Quark` seems to be a better *joint* ordering than can be obtained by the strategy of `SCALCE` (which imposes the reference free ordering derived for one of the reads on all of the mates) or `Mince` (which concatenates the read pairs and determines an ordering for the merged reads). Though `Quark` orders the reads according to the island to which the reads map (and the offset within that island), the fact that both reads are accounted for, simultaneously, when generating the mapping, seems to result in a superior joint ordering.

For SRR037452, the mapping rate is relatively low, which leads to worse compression rates for `Quark`, although `Mince` is the only tool that generates better compression than `Quark` (and only by a negligible margin). As unmapped reads are sent to `Mince` for compression, this comparable compression rate for datasets with low mapability is understandable.

## 5 Conclusion and Future Work

We have introduced `Quark`, a *semi-reference*-based method for the compression of RNA-seq reads. `Quark` uses a lightweight replacement for alignment, called quasi-mapping, which enables it to quickly find shared sequence among the input reads by determining the manner in which they were generated from the underlying reference. This structure is encoded in terms of the fragment equivalence classes and, eventually, the transcript islands that `Quark` uses for encoding. By taking advantage of this information, `Quark` is able to obtain compression rates better than existing tools aimed at the compression of raw sequencing reads. At the same time, `Quark` remains reference-free from the perspective of the decoder. This means that the decoding end is agnostic to the reference used to encode the reads, avoiding the fragility associated with encoding the reads with respect to a specific reference version, and allowing recovery of the sequencing reads from the `Quark` files alone.

While `Quark` already exhibits state-of-the-art compression performance, we believe that the opportunity exists for further improvements. For example, when reads are encoded with respect to the reference sequence, true divergence between the reference and the sample being encoded will be recorded multiple times (once for each read overlapping the variant). However, the same process that is used to encode the reads can be used to note and potentially detect the variant, as well as to distinguish between differences that result from random sequencing errors and those that result from true genomic variation. Thus, one could imagine that `Quark` could be used, simultaneously, to compress reads, detectnovelvariants, andtocorrectreaderrors. Though technically “lossy”, correction of the errors would be done in a fashion that would not change the mapping locations of the reads, and it would likely result in even better compression rates, as the random sequencing artifacts, which increase the entropy of the encoded stream, could be largely removed. Moreover, by applying the discovered variants to the transcript islands used for read encoding, the differences between the encoded reads and the reference sequence could be reduced even further, leading to more gains in sequence compression. This difference could be especially large when encoding is done with respect to atypical samples (e.g. cancer) or when encoding with respect to a *de novo* assembly.

Finally, the islands concept introduced in `Quark` opens up avenues for a data-driven approach to discovering novel splice junctions for non model organisms. The islands capture information about exons or co-spliced groups of exons. The segmentation of already assembled transcripts within equivalence classes can be characterized as a fine grained notion of previously unknown exonic regions.

